# Single-cell multimodal modeling with deep parametric inference

**DOI:** 10.1101/2022.04.04.486878

**Authors:** Huan Hu

**Affiliations:** Department of Physics, and Fujian Provincial Key Laboratory for Soft Functional Materials Research, Xiamen University, Xiamen, 361005, China; National Institute for Data Science in Health and Medicine, and State Key Laboratory of Cellular Stress Biology, Innovation Center for Cell Signaling Network, Xiamen University, Xiamen, 361005 China; Wenzhou Institute, University of Chinese Academy of Sciences, and Oujiang Laboratory (Zhejiang Lab for Regenerative Medicine, Vision and Brain Health), Wenzhou, Zhejiang 325001, China

## Abstract

The paired measurement of multiple modalities, known as the multimodal analysis, is an exciting frontier for connecting single-cell genomics with epitopes and functions. Mapping of transcriptomes in single-cells and the integration with cell phenotypes enable a better understanding of cellular states. However, assembling these paired omics into a unified representation of the cellular state remains challenging with the unique technical characteristics of each measurement. In this study, we built a deep parameter inference model (DPI) based on the properties of single-cell multimodal data. DPI is a complete single-cell multimodal omics analysis framework, which has built in multimodal data preprocessing, multimodal data integration, multimodal data reconstruction, reference and query, disturbance prediction and other analysis functions.

## Introduction

In recent years, advances in high-throughput single-cell multimodal measurements have continued to expand our understanding of cellular ontology, state, and function [1, 2]. Recently, the single-cell multimodal experimental techniques motivated by CITE-Seq [3] and REAP-Seq [4] have been developed to simultaneously measure the abundance of cell surface proteins, also called epitope, by extending the single-cell RNA sequencing (scRNA-seq) technology. Single-cell multimodal experimental data require synthetical methods for analysis with a complete cell view [5, 6].

Several state-of-the-art algorithms have been proposed to integrate single-cell transcriptome and epitope information, such as Seurat and totalVI. Seurat v3 was one of the first tools developed for single-cell multimodal analysis [7]. It performs the standard workflow of single-modality analysis of CITE-seq data to cluster cells, while examining these results using information from other modalities. However, it does not take the full advantage of multimodal data sourced from the same cells. As an upgradation, Seurat v4 introduces “Weighted Nearest Neighbor” analysis, which assigns weights to each modality in each cell, enabling a comprehensive analysis of multiple modalities [8]. TotalVI is an end-to-end federated analysis framework that probabilistically represents multimodal data as a combination of factors, thus eliminating the influence of technical factors on the integrated data [9].

In this study, we propose a deep parametric inference model (DPI), a new deep learning framework that integrates CITE-seq/REAP-seq data. With DPI, the cellular heterogeneity embedded in the single-cell multimodal omics can be comprehensively understood from multiple views. DPI is a complete single-cell multimodal omics analysis framework, which has built in multimodal data preprocessing, multimodal data integration, multimodal data reconstruction, reference and query, disturbance prediction and other analysis functions.

## Results

### DPI analysis framework

The workflow of DPI is shown in Fig. 1. First, single cell transcriptome and protein data are preprocessed separately. Single-cell transcriptome data are reported to obey binomial distribution or zero-inflated binomial distribution while protein data follow Poisson distribution[10–12]. We customized preprocessing strategies for both types of data. Next, transcriptome and protein data were fed into their respective encoders. They are transformed into low-dimensional data that fit a standard normal distribution. During this process, the mean and standard deviation parameters of their standard normal distributions are inferred by the neural network. The low-dimensional latent space constructed by the mean and standard deviation characterizes the properties of a single omics data. We separately mixed the mean and standard deviation of the two omics to infer the multimodal mean and standard deviation. The multimodal low-dimensional representation is constructed from the mean and standard deviation, which is also constrained by the standard normal distribution. Transcriptome, protein and multimodal latent spaces map cellular heterogeneity on a continuous standard normal distribution. These low-dimensional representations can be used for downstream analysis, where the multimodal space is the most complete, which can perform cell clustering, visualization, reference and query, and even prediction of the effects of genetic perturbations on cell states. Transcriptome and protein spaces are used to infer the parameters of the raw data distribution to restore ideal transcriptome and protein data patterns. In conclusion, we have developed a complete pipeline to integrate and analyze single-cell multimodal data.

**Fig. 1.**
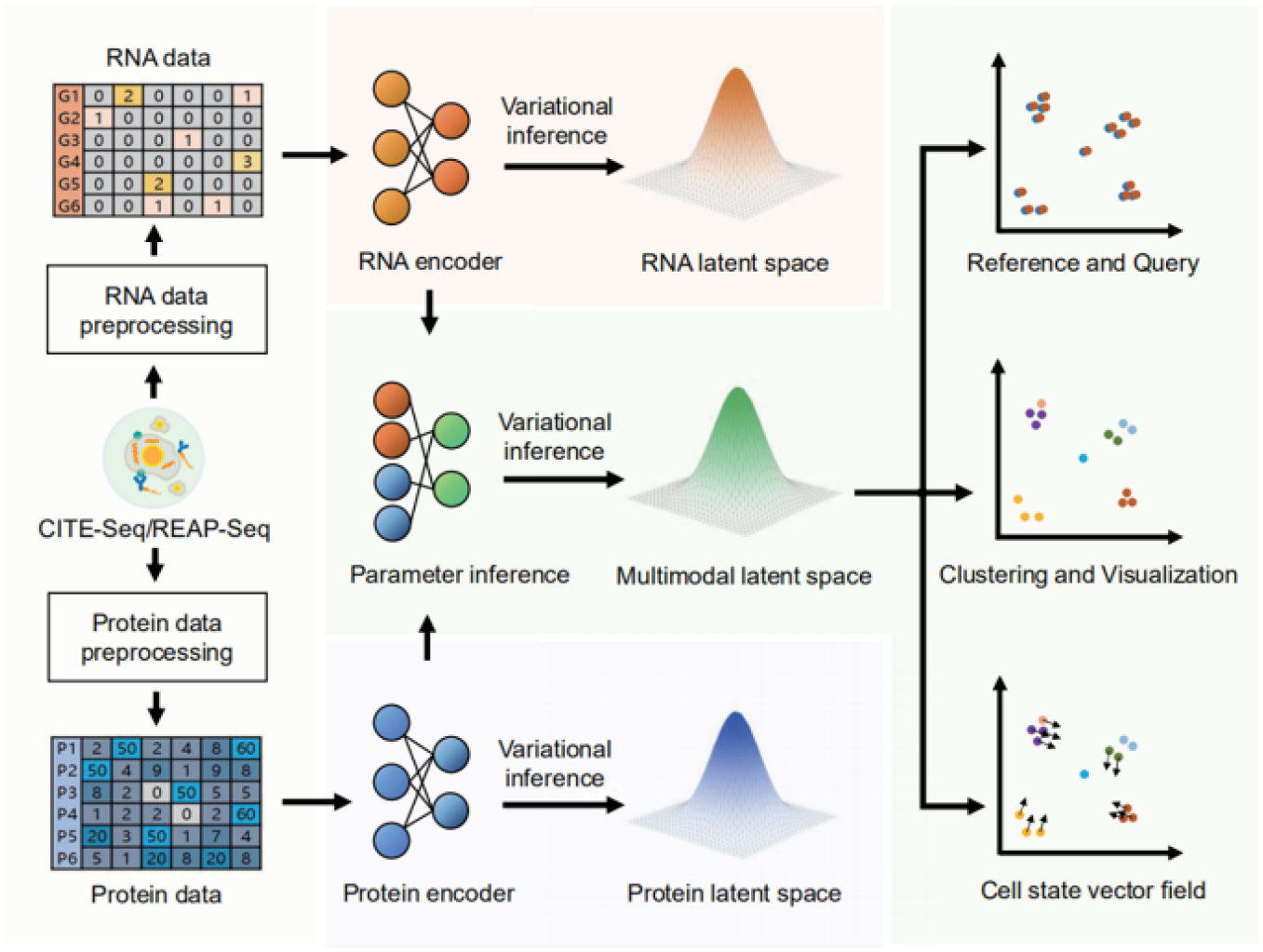
Workflow of DPI Analysis Framework.

### DPI enables efficient analysis of single-cell multimodal data

We performed DPI on the Cord Blood Mononuclear Cells (CBMC) dataset to reveal cellular heterogeneity. The CBMC dataset is derived from the earliest CITE-seq experiments [3]. It contains 8,617 cells and simultaneously measures 24,370 type genes and 13 type cell surface proteins. We performed DPI on the CBMC dataset and extracted multimodal low-dimensional representations, which were used for clustering and visualization. Fig. 2A shows the cell annotation for the CBMC dataset. We identified 14 cell subtypes on the CBMC dataset, and their corresponding markers can be found in Fig. 2B. The results show that the multimodal low-dimensional representation constructed by DPI can characterize the cellular heterogeneity of CBMC datasets. In addition, DPI restored the pattern of transcriptome and protein data of CBMC, respectively. Due to technical flaws, single-cell transcriptome sequencing can result in a degree of data loss, known as “Dropout”. As shown in Fig. 2C, DPI restored the CD14 gene distribution, which was closer to CD14 at the protein level than the original CD14 gene. Another example is the CD4 gene. The distribution of the DPI restored CD4 gene was in good agreement with the original CD4 gene distribution and was only specifically expressed on CD4+ T cells. Raw protein data may contain some noise, which is usually caused by the non-specific binding of the antibody tag [3]. As shown in Fig. 2 D, CD14 and CD4 that were restored by DPI were at lower levels in the cell clusters in which they were specifically expressed compared to the raw data.

**Fig. 2.**
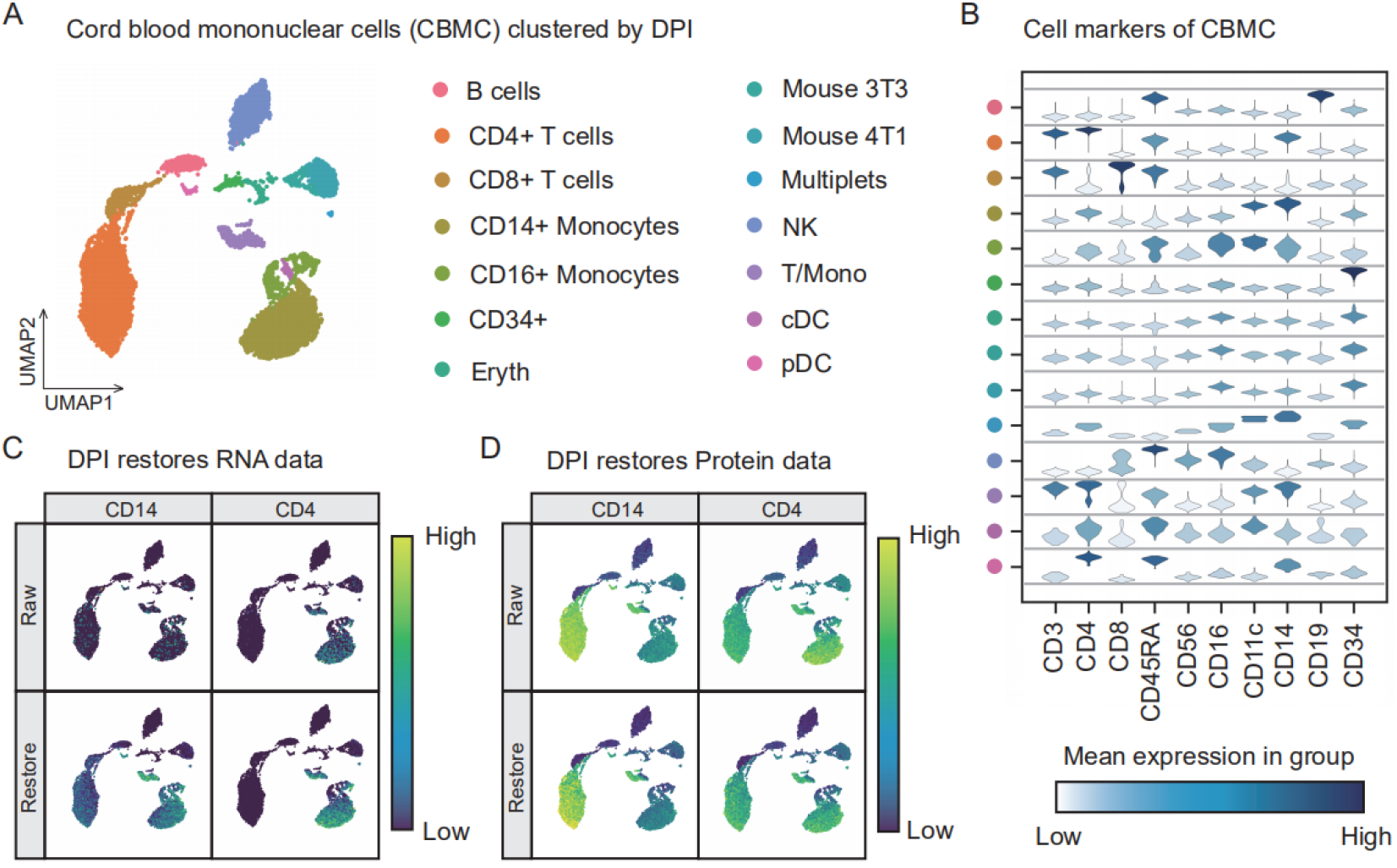
Applying DPI to analyze the CBMC dataset. The CBMC dataset is annotated with 14 cell subtypes (**A**) according to their corresponding markers (**B**). DPI also restored the distribution of RNA (**C**) and protein (**D**) simultaneously.

DPI can not only reveal the heterogeneity of cells but also have efficient performance. We compared DPI with SeuratV4 and TotalVI, which are standout models for single-cell multimodal analysis. The PBMC5K, PBMC10K and MALT10k datasets were involved in the evaluation of model performance, which contain 5,247, 7,865 and 8,412 cells, respectively. We evaluated the clustering results of the low-dimensional representations output by each model based on these three datasets. It is worth noting that to eliminate differences in clustering algorithms, the low-dimensional representations of these models are all fed into the Leiden clustering algorithm [13] with the same parameters. The results show that the DPI model outperforms SeuratV4 and TotalVI on both Calinski Harabasz score [14] and Silhouette score [15] evaluations (Fig. 3A). Finally, we tested the running time of DPI, SeuratV4 and TotalVI on different data scales under the same hardware environment. With the increase of the data size, the increase of the running time of the DPI model is smaller than that of SeuratV4 and TotalVI. This implies that the DPI model has better performance in the big data context.

**Fig. 3.**
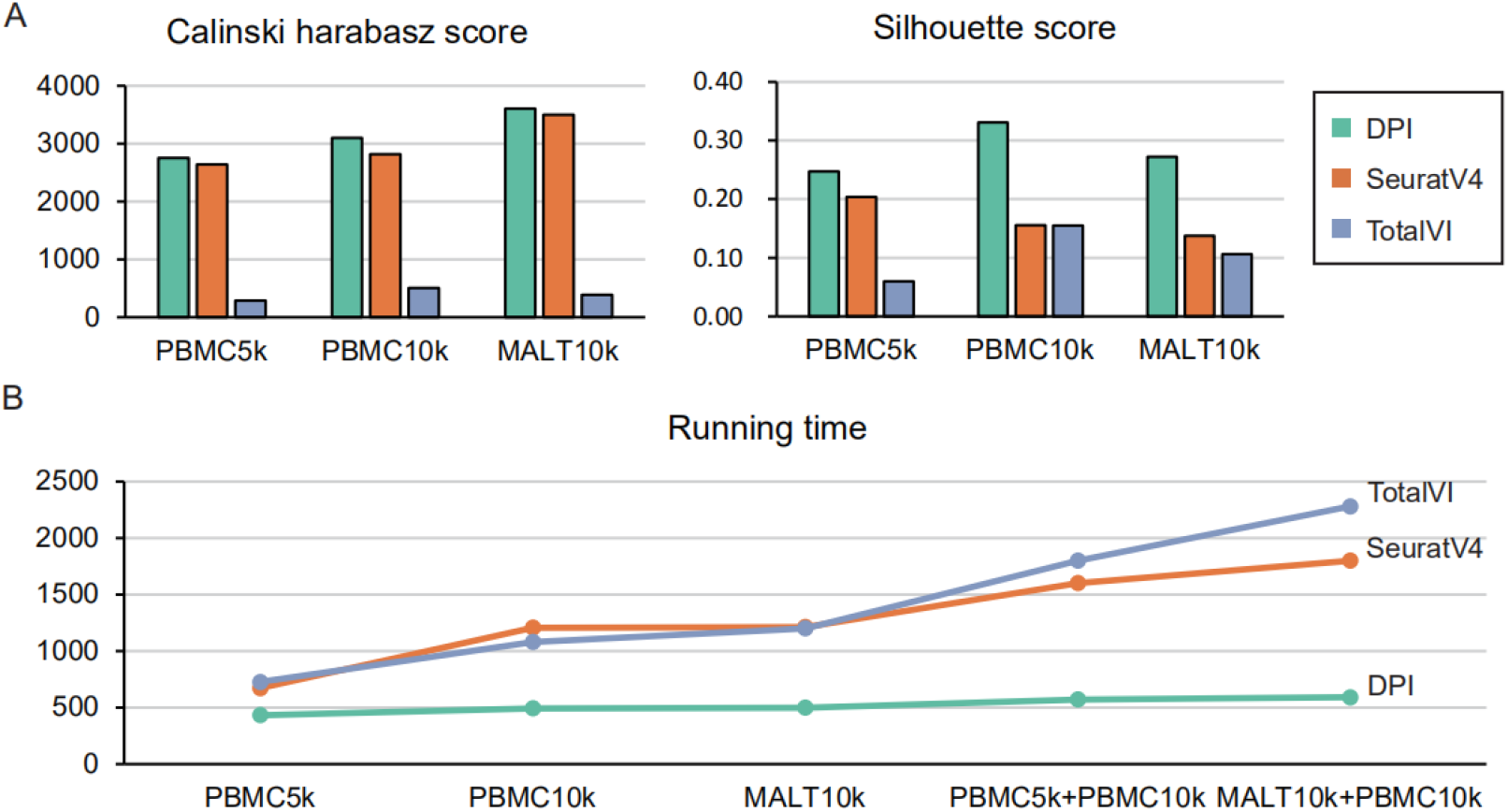
Performance comparison of DPI, SeuratV4 and TotalVI. We evaluate the performance of DPI, SeuratV4 and TotalVI based on two common clustering evaluation metrics (**A**) and running time (**B**).

### Reference and query of multimodal latent space

Raw single-cell data are limited. The multimodal latent space generated by DPI is a continuous normally distributed space, which is equivalent to a collection containing infinite single-cell data. When the raw multimodal data is transformed into a determined latent space by the DPI model, it can be used as a reference for unknown data. As shown in Fig. 4A, we clustered and visualized PBMC10K by the DPI framework. At this point a multimodal latent space representing PBMC10k has been established. Next, we feed the raw PBMC5k data into the PBMC10k trained DPI framework, which will be mapped in the multimodal latent space of PBMC10k to generate its low-dimensional representation. We project the low-dimensional representation of PBMC5k to the UMAP visualization of PBMC10k and find that they are aligned [16, 17]. This may be caused by the fact that PBMC10k and PBMC5k belong to the same tissue, and their mapping distances on the multimodal latent space are very close. Further, we queried PBMC5k for possible cell types based on its location in the PBMC10k latent space. The results of UMAP visualization suggest that the multimodal latent space can be used to query cell types (Fig. 4B).

**Fig. 4.**
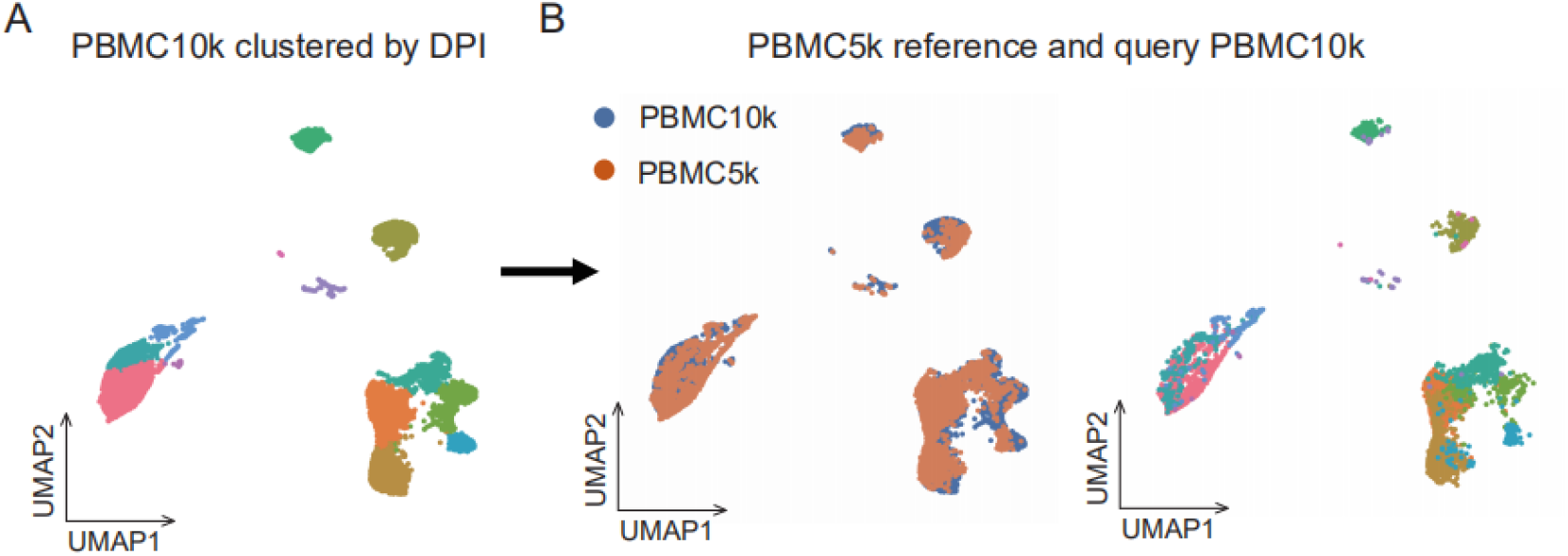
Apply the multimodal latent space of PBMC10k (A) to reference and query PBMC5k (B).

### Perturbation of Multimodal Latent Space

The above results have shown that DPI describes the state of all cells in the sample in terms of the multimodal latent space generated by the sample. Further, the multimodal latent space generated by DPI is continuous, which means that perturbing the genes/proteins of cells in the sample can find the cell state closest to it in this space. Taking the cell state before perturbation as the starting point and the cell state after perturbation as the end point, a vector field is established, which can describe the change of cell state. Fig. 5A shows how DPI perturbs multimodal data (using genes as an example) to generate vector fields describing cell state changes. First, perturbed multimodal data is fed into a pretrained DPI model. Next, the DPI model produces a perturbed multimodal latent space. Finally, the original multimodal latent space and the perturbed multimodal latent space are mapped to UMAP visualization and plotting cell vectors.

**Fig. 5.**
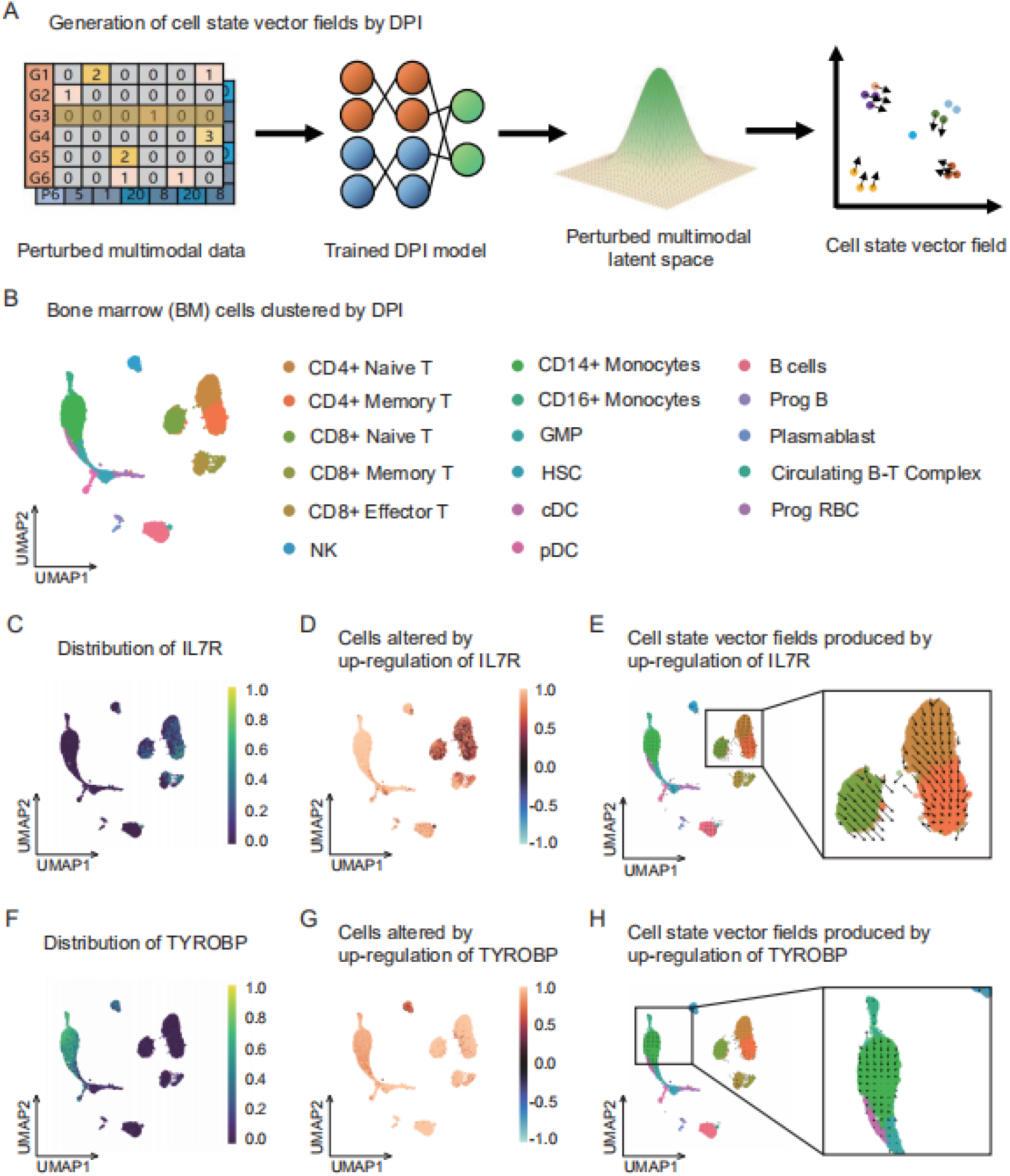
Applying DPI perturbation to a multimodal latent space to predict cell states. The pipeline of DPI to generate cell state vector fields (**A**) was performed on the BM dataset (**B**), and it successfully predicted changes in cell state upon upregulation of IL7R (**C,D,E**) and TYROBP (**F,G,H**).

To illustrate the function of DPI perturbation of the multimodal latent space, we analyzed bone marrow (BM) cells. Fig. 5B shows the cell types covered by the BM dataset. In BM samples, T cells had both native and memory states. Upregulation of the IL7R gene leads to a T cell state closer to a memory state. Fig. 5C shows the distribution of the IL7R gene, which is specifically distributed in the CD4+ T cell cluster. We up-regulated IL7R gene expression in BM samples to generate changes in the multimodal latent space. The difference between the multimodal latent space after IL7R upregulation and the original multimodal latent space was visualized by UMAP (Fig. 5D). We noticed that the cellular state of most cell clusters was not affected by upregulation of the IL7R gene except for CD4+ native T and CD4+ memory T. The state of CD4+ native T was biased towards CD4+ memory T after IL7R upregulation by observing the cell state vector field (Fig. 5E). This implies that upregulation of IL7R leads to the transition of T cells to a memory state, which is consistent with previous studies [18]. Another example is about monocytes. The TYROBP gene is highly expressed in mature monocytes [19]. Fig. 5F shows the distribution of TYROBP gene in cell clusters. After we up-regulated the TYROBY gene, there was a change in the cellular state of the cell cluster where the monocytes were located (Fig. 5G). Through the observation of the cellular vector field (Fig. 5H), we found that the states of monocytes did not shift outward, instead they were concentrated inside the monocyte clusters. This implies that high expression of TYROBP does not promote the transformation of monocytes to other cellular states but only to maturation.

## Discussion and Conclusion

DPI is a powerful single-cell multimodal data analysis framework, which can not only complete the integration and visualization of multimodal data, but also infer unknown cell types (reference and query) and predict cell state changes after perturbation. The success of DPI mainly stems from the multimodal latent space generated by the DPI framework. The multimodal latent space is the complete continuous whole inferred from the discrete multimodal data. To obtain the multimodal latent space, we extract the parameters of the single-model latent space for fusion. Considering generality and computational convenience, the standard normal distribution was chosen to build the multimodal and single-model latent spaces. The performance of a multimodal latent space depends on the ability of a single modality to capture cellular heterogeneity. In this study, we ensured that the single model accurately captures cellular heterogeneity information by decoding the single model space to the original data distribution space. DPI can also be used for other multimodal research projects (such as the integration of image and text information), and it has a wide range of applications.

## Method

### Quality Control

Quality control is a type of data screening strategy whose goal is to screen data to contain only high-quality real cells and features, so that when we perform downstream analysis, it is easier to capture reliable information.

For the transcriptome, three strategies were performed to screen for reliable gene expression data. First, each cell with total gene expression less than 200 in the transcriptome count matrix was deleted. Second, each gene was required to be expressed in at least 3 cells. Finally, cells with mitochondrial genes exceeding 20% of the total number of genes per cell in the count matrix were deleted. A high proportion of mitochondrial genes indicates poor cell quality, possibly due to loss of cytoplasmic RNA from perforated cells [20, 21]. The reason is that mitochondria are larger than individual transcript molecules and are less likely to escape through tears in the cell membrane. Generally, protein data, considering their few categories, are not screened unless they are experimentally proven to be contaminated.

### Normalization

In single-cell sequencing experiments, it is difficult to ensure consistency in library preparation between samples due to technical differences in amplification and capture efficiencies. This also causes systematic differences in the sequencing data of multiple samples due to differences in library sequencing coverage. The purpose of data normalization is to eliminate systematic differences while preserving data heterogeneity.

In this study, in order to maximize the heterogeneity of the transcriptome, the transcriptome data was logarithmically transformed, which has been widely used in single-cell transcriptome data preprocessing.

In this study, in order to maximize the heterogeneity of the transcriptome, the transcriptome data was logarithmically transformed, which has been widely used in single-cell transcriptome data preprocessing [12]. Previous studies have shown that the gene expression of each log-transformed single cell obeys a binomial distribution.

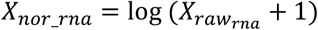

*X_raw_rna_* represents the raw single-cell transcriptome count matrix. *X_nor_rna_* represents the normalized transcriptome technology matrix. Considering the sparsity of transcriptome data (the expression of most genes is zero), *X_raw_rna_* is added with 1 to correct the possible negative infinity of logarithmic transformation.

Protein counts are treated as part of the whole cell, which is treated as component data and a centered log ratio (CLR) transformation is applied [22].

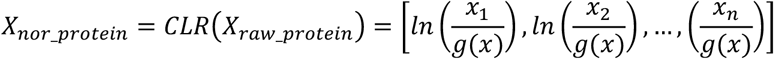

*Xraw_protein* represents the raw protein count matrix. *x* is a vector of protein counts (including one pseudo-count for each component) and *g*(*x*) is the geometric mean of *x*. We consider each type of protein in *X_nor_protein_* to follow a Poisson distribution.

### Feature selection

Transcriptome data feature selection methods refer to Seurat v3. We calculated and normalized the variance for each gene in the single-cell transcriptome count matrix. First perform z-score normalization on the data. Then, calculate the normalized variance for each gene. Finally, the genes are sorted to obtain highly variable genes. In fact, we call the “highly_variable_genes” function of the Scanpy python package (version 1.8.2) to perform feature selection on transcriptome data.

No feature selection was performed on the protein data considering that the feature dimensionality of the protein data is low and low-quality proteins have been removed in the experiments.

### Scaling

Transcriptome and protein data have different dimensions, which affect the results of data analysis. To eliminate dimensional effects between metrics, data scaling is required. Transcriptome and protein data were scaled to a range of 0 to 1 by *MinMaxScaler*, respectively, before being fed into the DPI model [23].

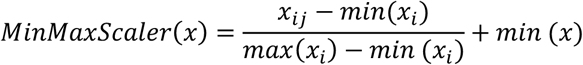

*x_ij_* represents the count of the *i-th* feature of the *j-th* cell. *min*(*x_i_*) represents the minimum value of the *i-th* feature. Similarly, *max*(*x_i_*) represents the maximum value of the *i-th* feature. *min*(*x*) is the minimum value in the entire cell count matrix, it is usually zero. *MinMaxScaler* preserves the distribution of the data while changing the data range.

*X_nor_rna_* and *X_nor_protein_* are scaled by *MinMaxScaler* to *X_scaled_rna_* and *X_scaled_protein_*, respectively.

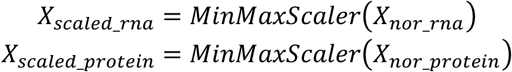

### Variational Autoencoder

Autoencoder (AE) is a neural network structure that encodes the input data first, and then reconstructs the original input only through the encoding [24]. Variational autoencoder (VAE) is a deep generative model based on the structure of AE, which describes the latent space *Z* and input samples *X* from a probabilistic perspective [25]. The goal of VAE is to get the distribution of *X* from data *X*. Since the distribution of *X* cannot be directly obtained, the latent variable *Z* is introduced, ie.

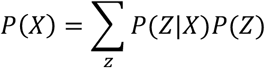

VAE encodes samples as probability distribution *P*(*Z*|*X*). Typically, *P*(*Z*|*X*) follows an isotropic normal distribution. The sampled vector obtained by random sampling from the latent variable *Z* can be decoded back to the data *X*.

### The DPI models

The DPI model is inspired by VAE, which consists of three parts: RNA parameter inference network, protein parameter inference network and multimodal parameter inference network.

The structure of the RNA parameter inference network is shown below. Notably, it is not a symmetric structure like a variational autoencoder. The goal of the RNA parameter inference network is not to match the input to the output but to infer the parameters of the distribution of the input data to minimize the difference in the distribution of the input and output. Here we assume that the RNA data follow a negative binomial distribution (NB).

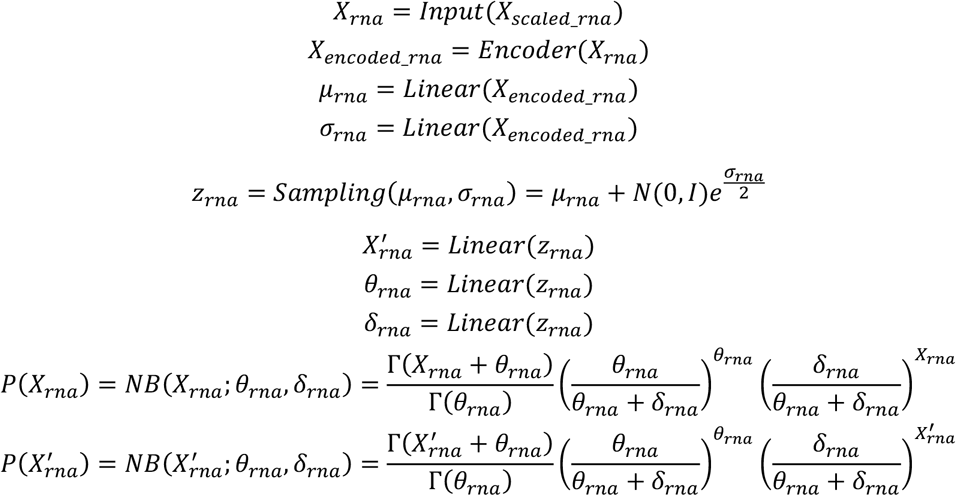

The structure of the protein parameter inference network is similar to that of the RNA parameter inference network. Here we assume that the protein data follow a Poisson distribution.

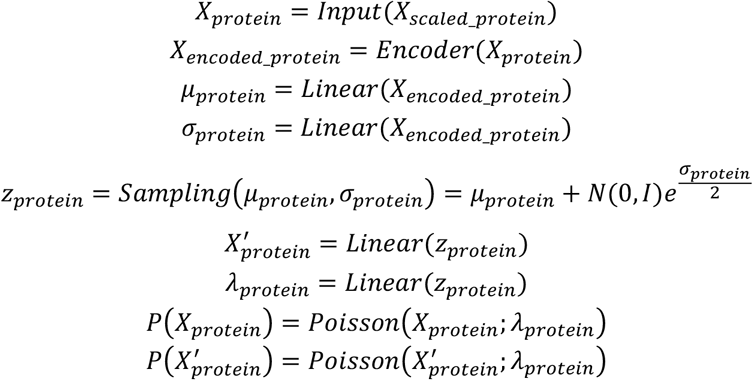

The multimodal parameter inference network relies on the RNA parameter inference network and the protein parameter inference network to provide parameters. Its goal is to restore the constructed multimodal standard normal distribution back to the standard normal distribution for RNA and the standard normal distribution for protein.

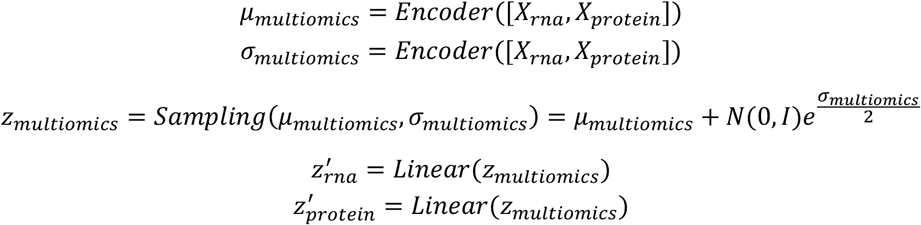

## Notes

### Competing Interest Statement

The authors have declared no competing interest.

